# Chronic Exposure to Palmitic Acid Downregulates AKT in Beta-Cells through Activation of mTOR

**DOI:** 10.1101/2020.08.11.247452

**Authors:** Richa Aggarwal, Zhechu Peng, Ni Zeng, Joshua Silva, Lina He, Jingyu Chen, Anketse Debebe, Eileen X. Stiles, Chien-Yu Chen, Bangyan L. Stiles

## Abstract

High circulating lipids occurring in obese individuals and insulin resistant patients are considered a contributing factor to Type 2 Diabetes (T2D). Exposure to high lipids initially causes the beta-cells to expand in population. Long-term exposure to high lipids however is associated with failure of beta-cells and the development of T2D. To prevent the failure of beta-cells and development of Type 2 Diabetes, this study focuses on understanding the molecular mechanisms that underlie this biphasic response of beta-cells to lipid exposure. Using palmitic acid (PA) in cultured beta-cells and islets, we demonstrated that chronic exposure to lipids leads to reduced viability and inhibition of cell cycle progression concurrent with downregulation of a pro-growth/survival kinase AKT, independent of glucose. This AKT downregulation by PA treatment is correlated with a consistent induction of mTOR/S6K activity concurrent with AKT downregulation. Inhibiting mTOR activity restores AKT activity and allows beta-cells to gain proliferation capacity that are lost after high fat diet exposure. In summary, we elucidated a novel mechanism for which lipid exposure may cause the dipole effects on beta-cell growth, where mTOR acts as a lipid sensor. These mechanisms can be novel targets for future therapeutic developments.

## Introduction

Macronutrients are recognized as factors that not only supply energy but also alter cellular growth response. These effects of glucose and lipids are particularly relevant to the function of pancreatic beta-cells as they respond to changes of plasma glucose and lipid levels to regulate insulin release. In individuals with normal blood glucose levels, this increased insulin requirement is met by increased insulin secretion from pancreatic beta-cells. In Diabetic patients, the beta-cells are unable to meet the increased insulin demands and patients manifest increasing glucose levels (hyperglycemia) and ultimately type II diabetes (T2D) (1–6).

Two factors contribute to the inability of beta-cell to compensate for the demand of hyperglycemia occurring in T2D: loss or reduction of the ability for beta-cells to secret insulin; and reduction of the number of beta-cells (2, 5, 7). Hyperglycemia and hyperlipidemia, the two conditions that hallmark insulin resistance, have both been hypothesized to regulate insulin secretion and the growth/survival of beta-cells (7). Glucose is known to stimulate beta-cell insulin secretion through regulation of the ATP-potassium transporter. It also stimulates the growth of beta-cells by regulating cyclin D2 expression (8, 9). The effects of lipids, however, are less clear compared to that of glucose. Depending on the type of lipid or duration of exposure, fatty acids can both stimulate or inhibit the function and growth/survival of cultured cells (10–14). Cultured human islets exposed to palmitic acid (PA) for 4 days led to apoptosis of beta-cells, whereas oleic acid protected against PA and glucose-induced beta-cell death (11). Similarly, prolonged exposure of rat or mouse islets to free fatty acids (FFAs) caused decreased insulin transcription, impaired glucose-induced insulin secretion (GSIS) and finally, resulted in beta-cell apoptosis (10, 15–18). However, increased beta-cell function has also been reported when islets or beta-cells are exposed to FFA (19–21). Rat islets treated with PA for 1 hour led to several fold higher BrdU incorporation versus controls concurrent with enhanced insulin secretion (19), indicative of enhanced cell growth and function. Overall, it appears that while short-term exposure of FFAs is pro-growth, long-term exposure is deleterious for the beta-cells.

In this study, we comprehensively evaluated the effect of short-term and chronic HFD feeding on islet mass and beta-cell proliferation. Our work demonstrated a clear biphasic effect of HFD feeding on beta-cell growth/survival. We then focused on the effects of saturated fatty acid PA on beta-cell growth and survival. Our results indicate that prolonged exposure of beta-cell to PA treatment caused suppressed proliferation and induced apoptosis, concurrent with inhibition of AKT activity, while short term exposure induced the pro-growth and pro-survival kinase AKT. We further elucidated mechanisms by which chronic exposure to PA suppresses AKT activity, and report a role of mTOR/Raptor mediated S6K activity in the regulation of AKT by PA.

## Results

### Chronic Exposure to High Fat Diet Leads to Reduced Beta-Cell Proliferation and Induced Beta-Cell Death

High Fat diet (HFD) is a lipid-rich diet that is often used as an experimental approach to induce hyperglycemia and insulin resistance conditions (4, 6). During HFD feeding, the mass of pancreatic islets increases to compensate the demand of insulin. It is hypothesized that this increased demand for insulin production leads to beta-cell death and development of diabetes. We used this HFD-induced compensatory beta-cell growth response as a model to explore the *in vivo* effects of hyperlipidemia on cell growth and survival. We subjected 3 months old mice to HFD feeding for different durations ranging from 7 days to 14 months. We found that 7 or 15-day feeding of HFD does not significantly alter the plasma insulin levels (Fig 1A). Consistent with previous reports (12), HFD feeding for 2 months is sufficient to increase fasting plasma insulin. Insulin levels peaked at 4 months, reaching approximately 10-fold induction vs. the normal chow fed mice. This induction is significantly diminished with 14 month-HFD feeding where only a 3-fold induction is observed (Fig 1A left panel). The mass of islets also increased concurrent with the elevated plasma insulin levels. Starting at 2 months post the start of HFD feeding, approximately 2-fold induction of islet mass is observed, reaching significance (p<0.05) with 4 months feeding. Islet mass peaked at 9 months with an approximately 4-fold induction and started to decline with 14 months HFD feeding (Fig 1A middle panel).

**Figure 1.**
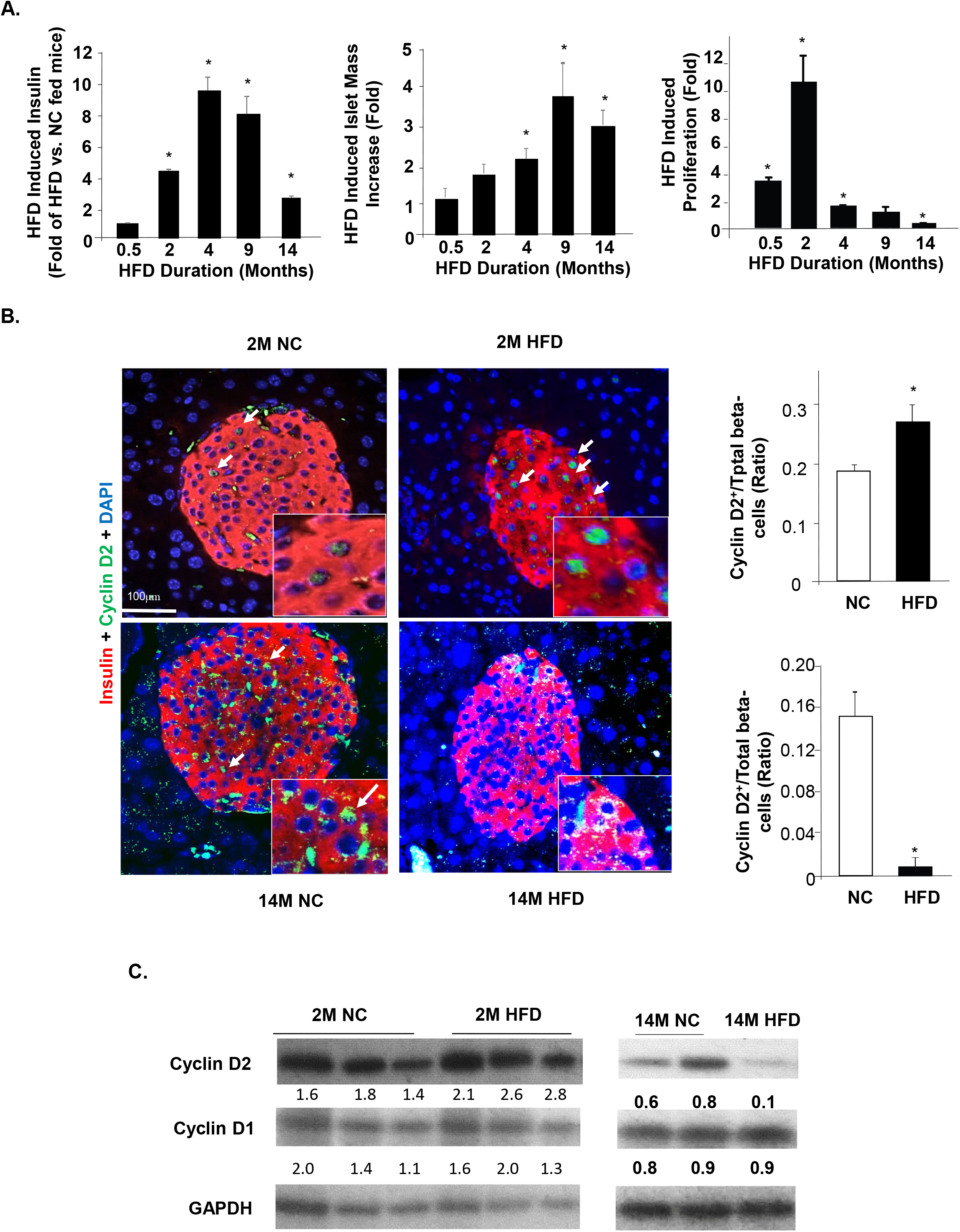
High fat diet induced beta-cell proliferation. **A.** 3 months old mice were put on high fat diet (HFD) for the indicated durations (X axes). Fold change of fasting insulin (left), Islet mass (middle) and cell proliferation rate (right) are reported for HFD vs. normal chow (NC) fed animals. n=5-9. *P<0.05. **B**. Immunofluorescent staining of cyclin D2 and insulin in pancreas of HFD and normal chow (NC) fed mice. Top panels, mice fed on indicated diet for 2 months (2M); Bottom panels, mice fed on indicated diet for 14 months (14M). Right panels, quantification of Cyclin D2+ beta cells vs. total beta cells. Green, cyclin D2; Red, Insulin; Blue, Dapi. n=5-9. *P<0.05. **C.** Immunoblotting analysis for cyclins D1 and D2. Left panel, mice fed on indicated diet for 2 months (2M); right panel, mice fed on indicated diet for 14 months (14M). Numbers under the row indicate ratio of the protein above over GAPDH.

We assessed the proliferation of beta-cells in response to HFD feeding. Our data suggests that increases in proliferation of beta-cell preceded the increase of either islet mass or plasma insulin concentration. Measured using Ki67/BrdU staining, our data showed that 14 day feeding of HFD is sufficient to induce beta-cell proliferation by 3 folds. Two-month HFD feeding induced beta-cell proliferation by nearly 10 fold (Fig 1A right panel). While longer feeding for 4 and 9 months continuously induced beta-cell proliferation, such induction occurs at approximately 2-fold or less, significantly lower than the 10-fold observed with 2 months HFD feeding. By 14 months of HFD feeding, beta-cell proliferation rate is significantly lower in the HFD fed mice. Approximately 50% beta-cell proliferation rate is observed in HFD feeding group vs. the control group. The cell proliferation rate is corroborated by cyclin D2 staining where 2-month HFD feeding significantly induced the number of cells stained positive for cyclin D2 (Fig 1B). In the 14-month-old mice, beta-cells positive for cyclin D2 are rare in the NC (Normal Chow/ Control) group and essentially undetectable in the HFD group (Fig 1B). Immunoblotting analysis of cyclin D2 confirms that 14-month HFD feeding led to reduced expression of cyclin D2 whereas 2-month HFD significantly increased levels of cyclin D2 (Fig 1C).

We also evaluated apoptosis in the beta-cells from the different HFD feeding groups using TUNEL analysis. In up to 9 months HFD feed mice, TUNEL positive cells are barely detectable in either normal chow or HFD groups (Fig 2 and data not shown). While TUNEL positive cells are still rare in the 14-month HFD fed mice, we were able to detect some TUNEL positive cells. In particular, some islets in the HFD group appear to be undergoing more severe apoptosis than the control group even though overall apoptotic rate is still very low (Fig 2). Nonetheless the increase in cell death in response to chronic lipid exposure likely contributed to the decline of insulin levels and islet mass associated with HFD feeding. Together, these *in vivo* analyses demonstrate a two-phased response of beta-cells to HFD feeding, where short exposure induces proliferation and overall islet function while longer exposure leads to apoptosis or cell death.

**Figure 2.**
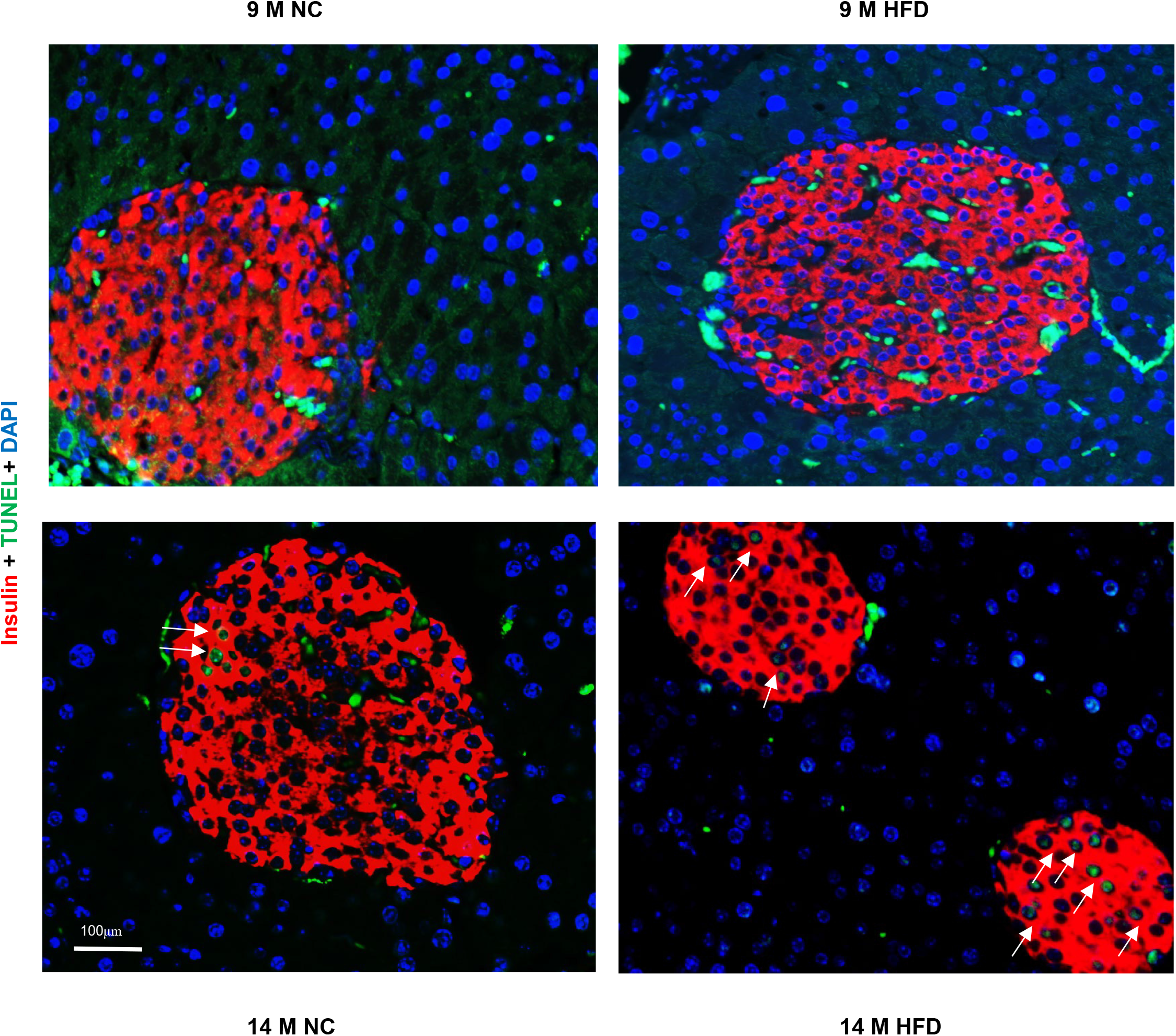
Chronic High fat diet induced beta-cell apoptosis. 3 months old mice were put on high fat diet (HFD) or normal chow (NC) for 9 and 14 months. Section of pancreas were stained with insulin (red) and TUNEL (Green). Blue, Dapi. Arrow: TUNEL positive cells.

### Prolonged Exposure to Palmitate Leads to Reduced Beta-Cell Growth/Survival

To understand this two-phased response to lipid exposure, we first performed time% and dose-dependent exposure studies to PA and evaluated cell viability using MTT assay in mEFs. We found that 0.4mM PA treatment can induce approximately 50% reduction in MTT in 24 hours while 48-hour treatment induced further reduction (Fig 3). Using the same criteria, we found that 48-hour treatment is needed for 0.4mM PA to inhibit MTT in beta-cells (INS-1, b-TC6 and MIN6) (Fig 3). Increasing concentration of PA dose dependently decreased cell growth/viability in mEFs as indicated by reduced MTT. In beta-cells, increased concentration of PA up to 1mM has minimum additional effects though a mild dosage effect is observed in INS-1 cells. Using the mEF cells as a model, we evaluated cell growth signals up to 24 hours with 0.4mM of PA treatment. Our data shows that PA indeed induced the expression of G1 cyclins, cyclin D2, similar to what we observed *in vivo* (Fig 4A). Eighteen and twenty-four hour treatments also induced the expression of cyclin D1 at physiological range of glucose. The induction of cyclin D2 by PA, however, appears to be independent of the glucose concentrations, though the specific exposure time needed does vary with different glucose concentrations.

**Figure 3.**
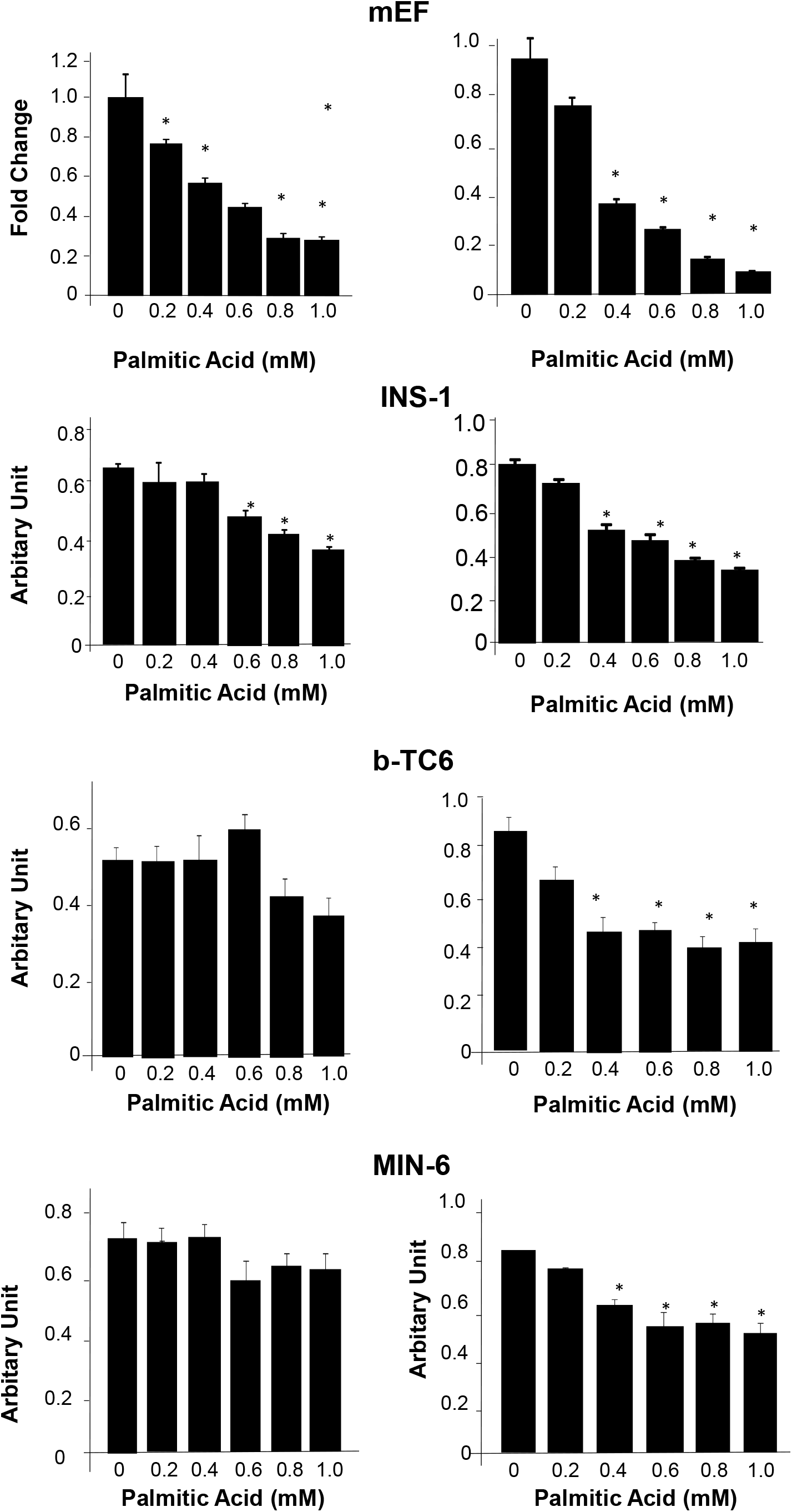
Prolonged exposure to palmitic acid treatment reduces cell survival/growth in multiple cell lines. In mouse embryonic fibroblasts (top panels) as well as three beta-cell cell lines (bottom three panels), exposure to palmitate acid for 24hrs (left panels) and 48 hrs (right panels) induced loss of cell viability/growth potential as measured with MTT assay. n=3. *P<0.05. Experiments repeated multiple times.

**Fig 4.**
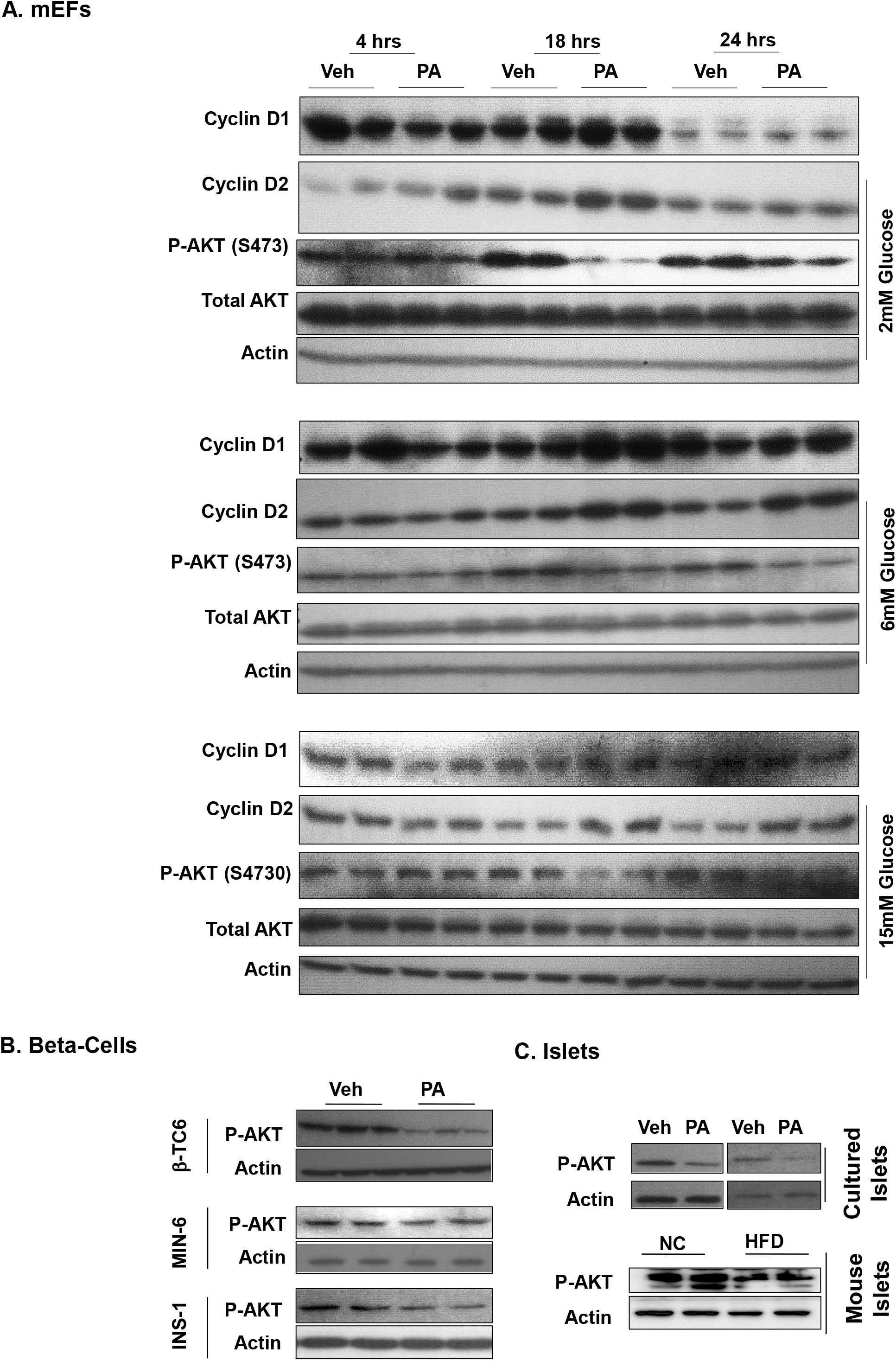
Chronic exposure to palmitic acid treatment induces cell cyclin D2 expression while inhibiting AKT phosphorylation. **A.** Mouse embryonic fibroblasts (mEFs) were exposed to 0.4 mM palmitic acid (PA) or DMSO (Veh) for indicated time points in the presence of indicated glucose concentrations. Cells were harvest and immunoblotting was performed for the indicated proteins. **B**. Three different beat-cell cell lines were exposed to 0.4 mM PA for 48 hours in the presence of 6 mM glucose. Immunoblotting for phospho-AKT confirmed downregulation of p-AKT by PA treatment. **C**. Top, isolated islets were cultured in the presence or absence of 0.4 mM PA for 72 hrs and immunoblotting for phospho-AKT showed downregulation of p-AKT by 0.4 mM PA treatment. Bottom, islets isolated from mice fed HFD for 4 months shows downregulation of p-AKT.

Surprisingly, we also observed a downregulation of phospho-AKT in all glucose conditions, particularly with 18- and 24-hour PA treatment. We confirmed this observation in the three beta-cell cell lines (Fig 4B). In addition, we exposed isolated islets to PA treatment and observed similar downregulation of AKT phosphorylation (Fig 4C, top panel). Furthermore, islets from HFD fed mice also displayed similar downregulation of AKT phosphorylation when compared with islets isolated from mice fed NC diet (Fig 4C, bottom panel). The downregulation of AKT phosphorylation is counterintuitive to the elevated G1 cyclin as AKT is a pro-growth and pro-survival kinase. We hypothesized that the downregulation of AKT maybe a result of chronic exposure to PA whereas short exposure would induce the phosphorylation of AKT. To address the dynamics of AKT phosphorylation in response to lipid exposure, we assessed AKT phosphorylation in INS-1 cells treated with 0.4mM PA for 30 min, 4 hour and 24 hours (Fig 5A). Our data clearly indicated that short treatment for 30 minutes indeed induced AKT phosphorylation whereas this phosphorylation is gradually lost with longer PA exposure. Twenty-four hour exposure to PA led to downregulation of p-AKT while 4 hour exposure does not alter the phosphorylation of AKT.

**Figure 5.**
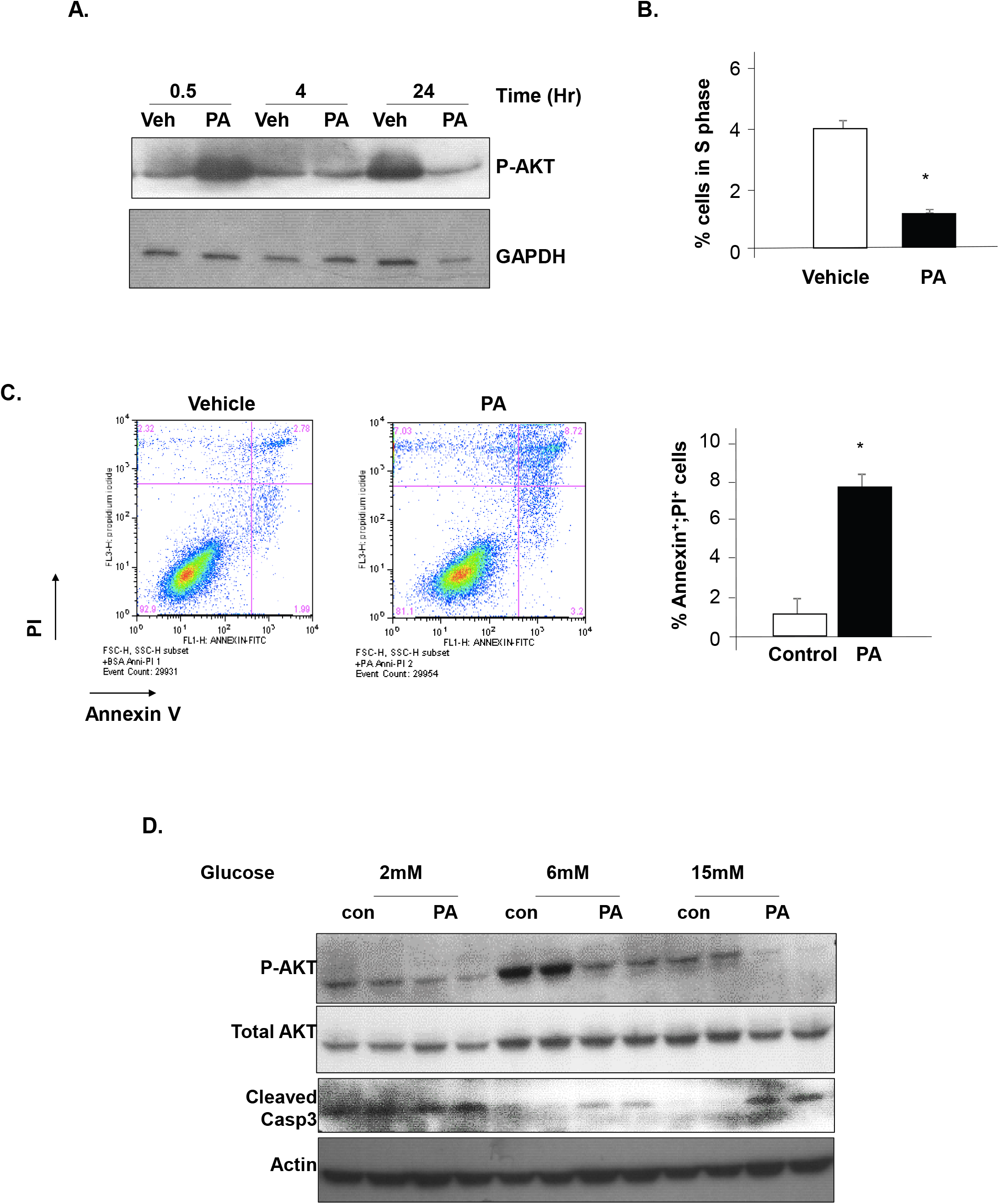
Biphasic response of phospho-AKT to palmitic acid and regulation of beta-cell growth and apoptosis. **A.** Short-term exposure to 0.4 mM PA reduced AKT phosphorylation while prolonged exposure induces it. INS-1 cells were exposed to 0.4 mM PA or DMSO (Veh) for the indicated time points. Immunoblotting is performed on lysates isolated from the cells for phospho-AKT. **B.** INS-1 cells were treated with 0.4 mM PA for 24 hours followed by flowcytometry analysis for the cell cycle. Percentage of cells in S phase is reported here. **C.** INS-1 cells treated with 0.4 mM PA for 24 hours were stained with annexin V and PI and analyzed using flowcytometry (left two panels). Percentage of cells dual positive for annexin V and PI are reported in the right panel. **D.** INS-1 cells treated with 0.4mM PA or DMSO (Veh) in the presence of different amount of glucose were lysed and proteins blotted for p-AKT, AKT and cleaved caspase 3 with actin as loading control. n=3. *P<0.05. Experiments repeated multiple times.

We hypothesized that the downregulation of AKT in response to long PA exposure may have been responsible for the loss of proliferation and increased apoptosis associated with prolonged lipid exposure. Consistent with this hypothesis, we observed 4-fold less cells in S phase in the 48 hr PA treated INS-1 cells (Fig 5B), indicative of reduced cell cycle progression. Furthermore, a 5-fold more Annexin V positive cells are observed in the PA treated cultures (Fig 5C), suggesting elevated apoptosis when INS-1 cells were exposed to PA for 48hrs. In addition, cleaved caspase 3, is induced by PA treatment, correlates with downregulation of AKT phosphorylation (Fig 5D). Together, these data indicate that pro-longed exposure of beta-cells to PA leads to reduced cell cycle progression and induces cell death.

### Prolonged Palmitate Exposure induced pAKT Involves Activation of p70S6K and Raptor/mTOR complex

We showed previously that AKT1 plays an important role in the adaptive response of beta-cells to HFD (22). Our data here suggest that PA regulated cell growth and survival is dependent on AKT phosphorylation where suppressed p-AKT is concurrent with PA induced beta-cell apoptosis and growth suppression. To explore how PA exposure leads to the downregulation of AKT phosphorylation, we first determined the level of PTEN as beta-cell deletion of *Pten* results in improved beta-cells function and survival of beta-cells (23, 24). In either mEF or beta-cell cell lines, PTEN does not change in response to PA exposure (Fig 6A). In islets, PA treatment appears to have a minor effect on PTEN expression (Fig 6A). Thus, it is unlikely that the downregulation of AKT phosphorylation induced by PA is due to PTEN induction.

**Figure 6.**
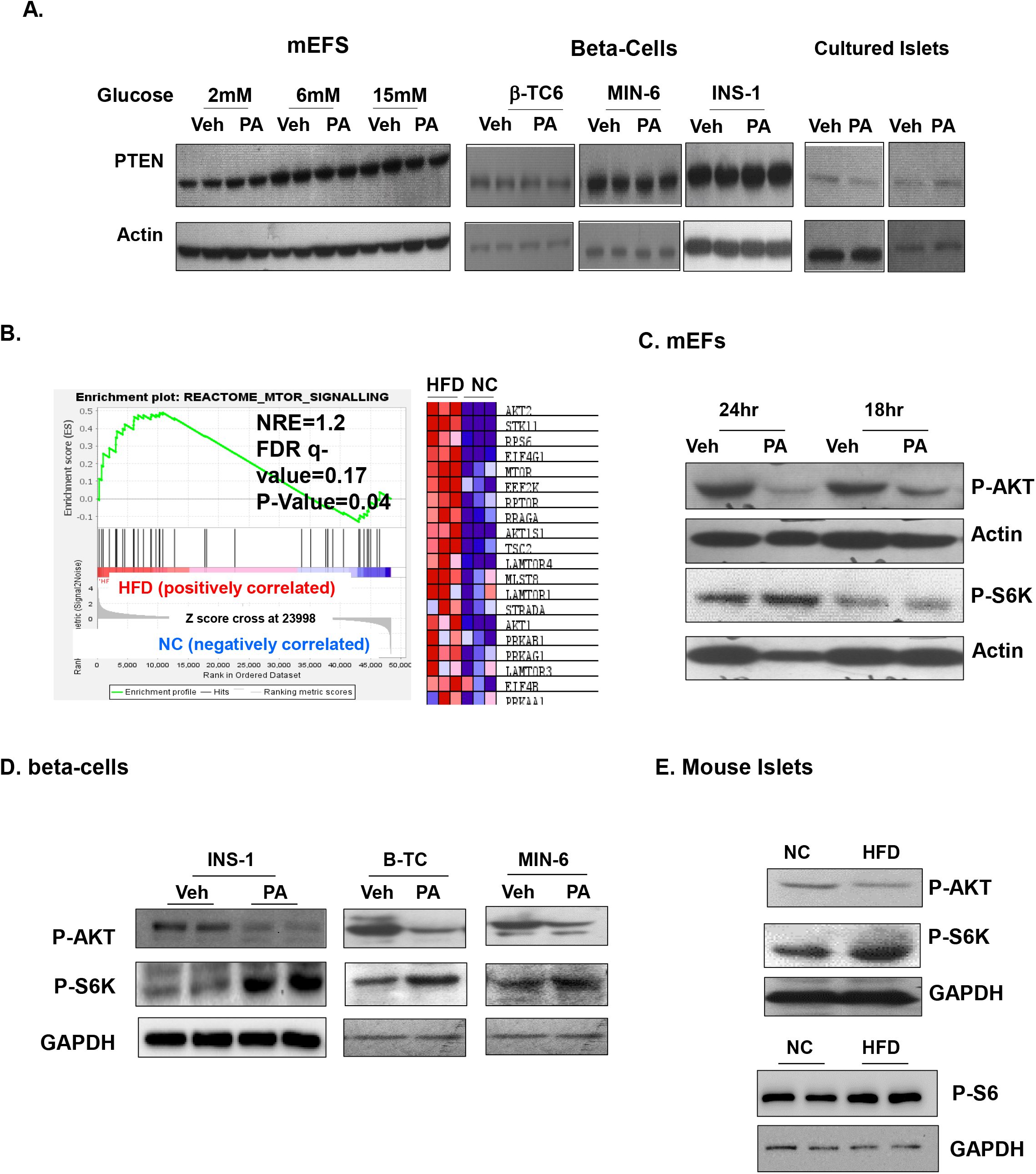
S6K signal but not PTEN is altered by exposure to palmitic acid. **A.** mEF, beta-cells, and cultured islets treated with 0.4 mM PA or DMSO (Veh) were lysed and analyzed for PTEN expression. **B.** Gene Enrichment Analysis of mTOR signal genes in islets from mice fed HFD for 4 months vs. those fed normal chow (NC). **C**. mEF cells exposed to 0.4 mM PA or vehicle with 6 mM glucose were analyzed for p-AKT and p-S6K. **D**. Three beta-cell lines were treated with 0.4mM PA with 6mM glucose and analyzed for p-AKT and p-S6K. **E.** Islets isolated from HFD fed mice (4 months HFD) were analyzed for p-AKT, p-S6K and p-S6.

In order to explore the mechanisms by which chronic HFD exposure lead to reduced p-AKT, we performed RNA-seq analysis in islets from HFD fed mice (submission to GEO database pending). Our data demonstrate that mTOR/p70S6K is among the top three major signaling pathway that is induced in islets by HFD feeding (supplemental Fig 1). Gene set enrichment analysis (GSEA) showed a significant upregulation of mTOR signaling in islets isolated from the HFD fed mice vs. controls (Fig 6B). To confirm that mTOR signal is indeed induced even with downregulation of AKT activity, we determined S6K phosphorylation in response to PA treatment. In mEFs exposed to PA for 24 and 18 hours, we found that PA treatment induced phospho-S6K even though phospho-AKT is inhibited, particularly with 24 hours exposure (Fig 6C). The same association of phospho-AKT and phospho-S6K is observed with beta-cells exposed to PA for 48 hours (Fig 6D). We also observed similar induction of S6K phosphorylation as well as its target S6 in islets from mice fed HFD diet (Fig 6E). These data together suggest that prolonged lipid exposure, while inhibiting AKT activity is inducing the activity of mTOR.

While mTOR activation is often indicative of active AKT activity, it also serves as the kinase that phosphorylates AKT. In addition, chronic activation of mTOR signal is reported to induce a S6K dependent feedback loop to inhibit AKT activity (25). To address if the PA induced AKT downregulation is dependent on this feedback loop, we treated INS-1 cells with rapamycin in combination with PA. As an inhibitor for mTOR activity, rapamycin treatment effectively inhibited phosphorylation of S6K. Our data indicated that the PA induced downregulation of AKT phosphorylation is readily rescued by rapamycin treatment (Fig 7A). This data suggests that the PA regulated downregulation of phospho-AKT is dependent on mTOR signaling. To address how PA may regulate mTOR signal, we determined the expression of Raptor and Rictor. Our data indicates that the overall protein levels of Raptor (but not Rictor) is increased with PA treatment (Fig. 7B). As activation of the Raptor-containing mTOR complex leads to phosphorylation of S6K, this data is consistent with the notion that the upregulation of Raptor containing mTOR complex by PA treatment is responsible for the S6K dependent downregulation of AKT phosphorylation.

**Figure 7.**
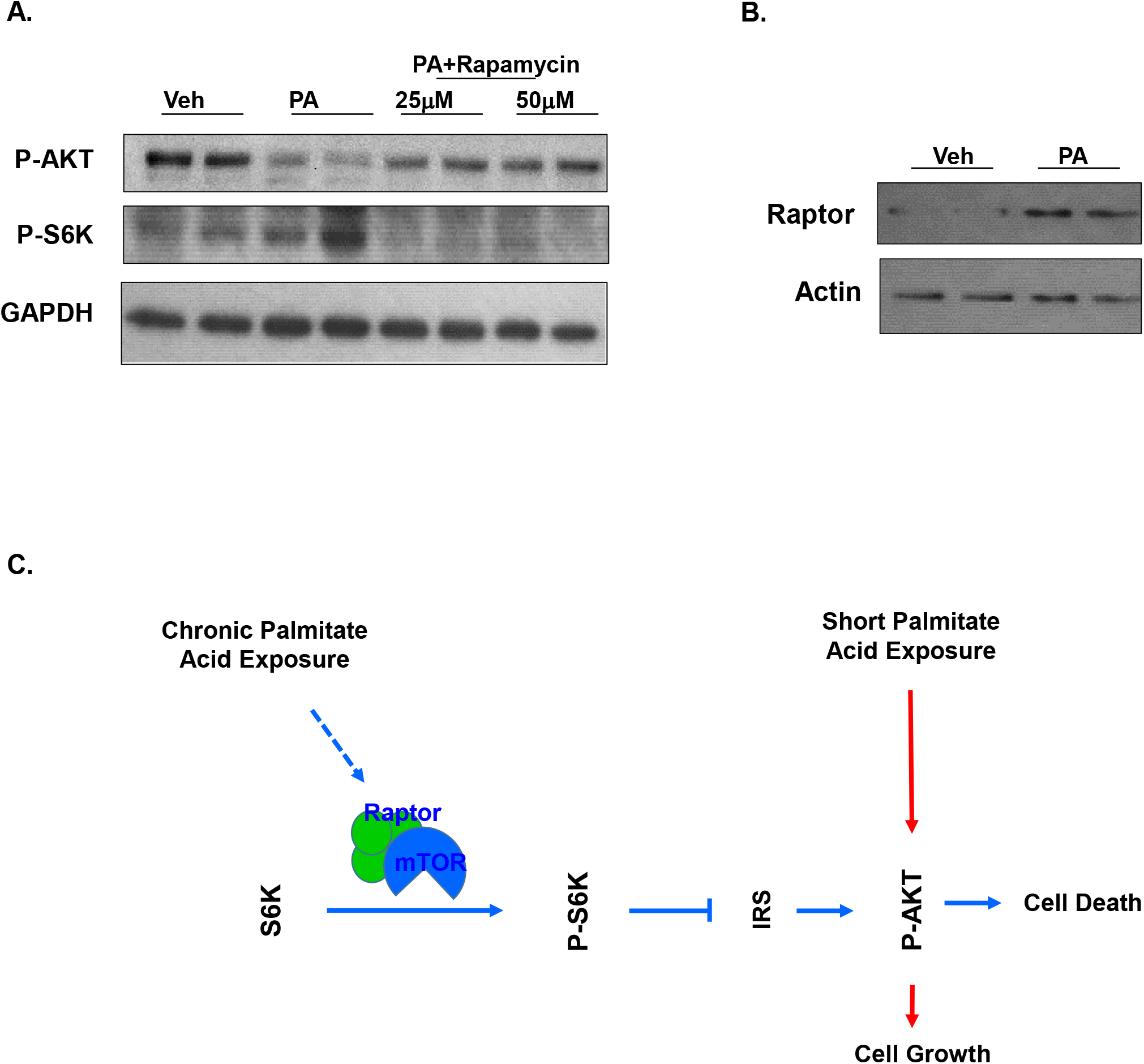
Induction of mTOR-S6K led to downregulation of AKT phosphorylation due to exposure to palmitic acid. **A**. INS-1 cells were treated with PA or PA+Rapamycin and analyzed for p-AKT and p-S6K. **B**. INS-1 cells treated with PA or vehicle were analyzed for Raptor. **C.** Proposed role of palmitic acid on loss of beta-cell function via S6K regulated AKT phosphorylation.

## Discussion

Obesity, a major complication associated with T2D, increases the demand of insulin secretion to cope with hyperglycemia pressure (3). The increased insulin secretion is accompanied by the increased proliferation of pancreatic beta-cells (2–6). This enhanced growth response from beta-cells is followed by failure of beta-cells where both the mass and function of beta-cells decline when the demand for increased insulin output sustains for chronic period of time (3, 26, 27). In recent years, the ability of glucose to induce beta-cell replication has been established (8, 28), however whether lipid induces beta-cell replication or promote their apoptosis is still unclear. Studies focusing on the role of lipid or HFD suggests that short term and low dose exposure to lipid is likely pro-growth for beta-cells whereas long-term and high dose exposure results in death of beta-cells (3, 11, 26, 29–32). However, the dynamics and mechanism by which lipid can induce both beta-cell growth and their apoptosis is not clear. Our study explored the dynamic response and mechanistic base for the lipid induced beta-cell compensatory response and reports several major findings. *First*, we comprehensively determined the dynamics of beta-cell response to HFD feeding. We show here that beta-cell proliferation peaked at 2 months after initiation of HFD feeding, preceding the peaks in increased insulin secretion (4 months) and islet mass (9 months). We also demonstrated that chronic exposure to HFD (14 months) resulted in reduction of beta-cell proliferation and elevated apoptosis. *Second*, we established cell culture models that mimic the bi-phasic response to chronic lipid exposure with multiple beta-cell cell lines. Using these model systems to confirm what was observed in mouse models, we discovered that chronic exposure (>4 hours) to PA leads to downregulation of phospho-AKT whereas short exposure (30 min) results in its induction. *Third*, downregulation of phospho-AKT occurs simultaneously with increases in annexin V positive apoptotic cells as well as reduction of cells in S phase. *Fourth*, downregulation of phospho-AKT by PA is not a result of altered expression of PTEN but regulated by Raptor-mTOR mediated S6K activation.

A “beta-cell exhaust” idea has been proposed where rapid growth of beta-cells induced by HFD leads to exhaustion of their growth capacity resulting in failure (33). Our study here started by exploring the molecular signals induced by HFD that regulate beta-cell growth. Using PA treatment in cultured beta-cells and islets, our data demonstrated that chronic exposure to lipids leads to reduced viability and inhibition of cell cycle progression concurrent with downregulation of a pro-growth/survival kinase AKT, independent of glucose. Genetic studies targeting AKT and signals regulating AKT have demonstrated a role for the AKT isoforms in the regulation of beta-cell growth/survival and islet mass (23, 24, 34–43). AKT2 was found to be required for maintaining metabolic homeostasis as *Akt2*^−/−^ mice develop insulin resistance, which indirectly leads to induced beta-cell mass due to adaptive beta-cell response (35). The role of AKT1 in metabolic regulation lies in its ability to regulate the adaptive growth and survival of pancreatic beta-cells (22). Mice deficient for AKT1 function display normal β cell mass and morphology (36, 41), whereas ectopic overexpression of constitutively active AKT1 in beta-cells leads to a dramatic increase in islet mass (34, 43). In addition, deletion of *Pten* in beta-cells which led to constitutive activation of AKT1 resulted in increased β cell proliferation, enhanced islet mass, and hypoglycemia in mice (23, 24). Consistently in beta-cells, AKT1 was found to be indispensable for the adaptive growth response for beta-cells fed HFD (22). Loss of AKT1 function induces mild ER stress and predisposes them chronic HFD induced cell death (22, 44).

In the past two decades, studies using genetically modified animals suggest a major role for the G1/S cell cycle machinery and key mitogenic signals such as IGF, PDGF and HGF in the growth of beta-cells (45). Among the molecular signals that control beta-cell replication, genetic evidence targeting different signaling molecules confirmed the importance of PI3K signal downstream of the growth factors in the regulation of beta-cell replication. The negative regulator of this mitogenic signaling, PTEN was previously shown to be induced by HFD feeding *in vivo* with unclear mechanisms (46). Our group reported that mice lacking PTEN specifically in the beta-cells have more and larger islets and demonstrated a role of PTEN/PI3K signaling in beta-cell growth and senescence (23, 38, 44, 47). Here, we report that the PA induced AKT downregulation is not a result of induced expression of its negative regulator PTEN but due to the feedback regulatory loop mediated by mTOR. Our data found that chronic exposure of beta-cells and islets to PA results in concurrent activation of S6K and inhibition of AKT. This analysis suggests that the mTOR-AKT negative feedback loop signaling is induced by chronic exposure to lipids and that this feedback signal is likely responsible for the dipole response of beta-cells to lipid/HFD exposure (Fig 7C). Though transient activation of mTOR leads to enhanced cell survival, chronic activation of mTOR results in inhibition of PI3K/AKT action via IRS and promotion of cell death (48). Such feedback can block the action of PI3K and results in the downregulation of downstream signaling molecules, including Ser/Thr kinase AKT (25). Consistently, mice lacking either TSC1 or TSC2 (with activated mTOR signal) in beta-cells display beta-cell failure with reduced islet mass and function and develop diabetes like phenotypes when they get older (49). Such failure is mediated by the activation of mTOR as rapamycin treatment to inhibit mTOR activity can rescue the failure of beta-cells in these mice (49). Interestingly, the rapamycin treatment also induced the phosphorylation of AKT.

The mTOR kinase exists in two separate complexes with other regulatory factors (48). Previous studies have reported that Myc dependent adaptive response to glucose is regulated by the TORC1 mTOR complex (50). The TORC1 complex is composed of Raptor and PRAS40. TORC1 complex phosphorylates and activates S6K which phosphorylates and inactivates IRS1/2. The TORC2 complex is composed of Rictor, mSin1 and Protor. Chronic activation of mTOR also leads to TORC2 induced phosphorylation and degradation of IRS1/2. Our data demonstrated a consistent activation of S6K phosphorylation in response to lipid treatment. These data suggest that lipid treatment at least activates the TORC1 complex, an observation confirmed by the upregulation of Raptor. In experimental models, loss of mTORC1 signal also lead to beta-cell failure and results in Diabetes phenotypes (42). However, inhibition of S6K led to improved GSIS in isolated human islets (51). Thus, further exploration into the mTOR-S6K-AKT is needed to understand the dynamic of response of this signaling axes to lipid exposure. Particularly, understanding the mechanisms by which lipid exposure regulates mTOR signaling is not only necessary for elucidating the adaptive response of beta-cells to lipid exposure, but also other cell growth responses to dyslipidemia. A putative mechanism characterized for lipid-mTOR interaction impinges on phosphatidyl acid (52, 53). Binding of phosphatidic acid to the FRB domain of mTOR blocks binding of Deptor (54), a partner of both TORC1 and TORC2 complex. While initial immunoprecipitation in INS-1 cells did not confirm that PA treatment altered binding of Deptor to TORCs (data not shown), our data shows that total levels of Raptor is induced by PA treatment in INS-1 cells. Raptor is an component of TORC1 complex and elevated Raptor is consistent with the observed increase of pS6K in response to PA treatment.

Together, our data suggests that Raptor-mTOR may act as a lipid sensor for HFD and increased lipid levels can induce beta-cell proliferation followed by beta-cell failure due to the mTOR feedback loop. Induction of Raptor expression and activation of S6K mediates downregulation of AKT due to chronic exposure to lipids. The downregulated AKT signal led to loss of growth potential and increased beta-cell death, leading to beta-cell failure in response to chronic lipid exposure. An increased activation of S6K has been recently reported in islets of human T2D patients where hyperlipidemia is common (51). In these islets from T2D patients and db/db mice, inhibition of mTORC-S6K signal can indeed improve GSIS (51). Our study here showed that the mTOR feedback loop mediates the dipole effect of HFD and lipid induced beta-cell growth deficiency and it may be targeted to overcome beta-cell failure.

## Experimental Procedures

### Animals

Mice of mixed backgrounds were used for the high fat diet (HFD) studies since different backgrounds have been shown to have different response to high fat diet. All animals were housed in a temperature-, humidity-, and light-controlled room (12-h light/dark cycle), allowing free access to food and water. All experiments were conducted according to the Institutional Animal Care and Use Committee of the University of Southern California research guidelines.

### Cell Culture

Various beta-cell (bTC6, MIN-6 and INS-1) and non-beta (mEFs) cell lines were used for the study. INS-1 cells were kindly supplied by Prasanna Dadi at Vanderbilt University. All cells were cultured at 37 °C in a humidified atmosphere containing 5% CO2 in either DMEM containing 25mM glucose or RPMI 1640 medium containing 11 mM glucose and supplemented with 10% heat-inactivated FBS, 50 μM 2-mercaptoethanol, 100 units/ml penicillin and 100 μg/ml streptomycin. Cells were starved overnight 48 hours after seeding and then treated with 0.4 mM PA in media containing 2, 6 or 15 mM glucose as indicated.

### Diet Feeding

For the high fat diet (HFD) experiment, one group of mice was fed with High Fat Diet with 60 kcal% fat (TD06414, Harlan laboratories) (55) whereas the control group was fed with Normal chow with 13 kcal% of fat in their diet (PicoLab 5053). Diet was started at 3 months of age and continued for the indicated period of time. Body weight, food intake and random plasma glucose levels were monitored weekly. BrdU (1mg/mL in deionized water, BD Pharmingen) was given to mice in water for 5 days before the ending of the study. Pancreas were collected at the end of the 5 days following overnight fasting.

### Biochemical analysis

Plasma samples were also collected through cardiac puncture at the end of the study following overnight fasting. Mouse Ultrasensitive Insulin ELISA kit from ALPCO (Cat#80-INSMSU-E01) is used for quantifying plasma Insulin per kit instructions as described previously (23, 24).

### Mouse islet isolation

Pancreases were perfused with collagenase P solution (0.8 mg/mL; 5mL per mouse) and digested at 37 °C for 17 min. Islets were then purified by using Ficoll gradients with densities of 1.108, 1.096, 1.069 and 1.037 (Cellgro) as previously reported (23, 24). Islets between layer 1.096 and 1.069 were collected and handpicked for either protein extraction or RNA extraction (26, 31).

### Rapamycin treatment in mice

Mice were fed with HFD for the required duration and rapamycin was injected intraperitoneally on the last 8 days of treatment (everyday 0.3mg/kg per mouse). 100mM Rapamycin stock in DMSO (LC labs # R-5000) was diluted 100 folds by mixing 890ul PBS, 100ul Tween 80 and 10uL rapamycin stock. Calculations were then done and mice were injected as per their body weight.

### Immunohistochemistry

At the end of the study, pancreas were dissected *en total* and fixed in Zn-formalin at 4^°^C for histopathology and immunohistochemistry analysis. Zn-formalin fixed and paraffin embedded sections were stained with hematoxylin and eosin for histopathology analysis (56). Islet mass is calculated based on area of islets vs. total pancreas area measurement collection from 3 sections per sample 60uM apart as described (23, 57).

Pancreas sections were also stained with antibodies for immunohistochemical or immunofluorescence analysis. Antibodies used are: Cyclin D2 (Santa Cruz, sc-593), Ki-67 monoclonal Ab (CST#12202) and BrdU (BD Pharmingen). TUNEL kit from BD Pharmingen were used for assessment of apoptotic cells following manufacturer’s instruction as described (47, 58).

### Fatty Acid Preparation

Palmitic Acid (sigma#P0500-10G) 200mM stock was prepared by dissolving 51.2mg PA in 1ml 100% ethanol. 0.04ml of this stock was further dissolved in 1.96ml 10% fatty acid free BSA (in DMEM media) for 4mM PA stock by shaking overnight at room temperature. On the following day, solution was filtered through 0.22μM filter and diluted 1:10 with media prior to cell treatment. 1% BSA in DMEM media was used as a control.

### MTT Assay

Cells were seeded at density 1.5-2×10^4^cell/well in 96-well plates in RPMI media. After 48 hours of seeding, the cells were either treated with 1% BSA (as control) or different concentrations of Palmitic Acid (0.2mM, 0.4mM, 0.6mM, 0.8mM and 1mM) (6 wells per treatment) and incubated for 24 or 48 hours. 10uL MTT reagent was then added after respective incubation and kept at 37^°^C for 4 hours, followed by addition of 100uL DMSO. The plate was kept on the shaker for 5-10 minutes to dissolve the crystals. Optical Density Reading was then taken at 570 nm wavelength.

### Cell Cycle FACS

INS-1 cells (3×10^5^ cells/well) were seeded in six-well plates in RPMI media with 6mM glucose with 10% FBS. Cells were starved for 48-72 hours in media without FBS after 24 hours of seeding. They were then treated with 1% BSA or 0.4mM Palmitic Acid for 48 hours. Cells were harvested by washing, trypsinizing and centrifuging them. The resulting pellet was re-suspended in 0.1mL PBS. 1mL ethanol was then added (kept at −20^°^C) and kept for 20 mins at −20^°^C. Cells were centrifuged at 1000rpm for 5 mins and supernatant was discarded. Then the pellet was resuspended in 500uL of RNase solution (200ug/mL in PBS) and incubated at room temperature for 30 mins. Propidium Iodide was later added to the mix (50ug/mL) and incubated for 30 mins away from light. FACS analysis was performed using FACS calibur machine and software.

### PI/ANNEXIN V FACS

INS-1 cells were seeded at density (2.5×10^5^ cells/well) in six-well plates in RPMI media. After 48 hours of seeding, cells were treated with 1% BSA or 0.4mM Palmitic Acid for 48 hours. Media containing detached and floating cells was collected and the rest of the cells were trypsinized, and washed once with PBS. Cells were then washed and suspended in 1X Annexin Binding Buffer (ABB). Diluted Annexin (400ng per 1×10^6^ cells) was then added to cells and incubated for 8 minutes at room temperature away from light. Propidium Iodide was then added to each sample (2.5ug/mL per sample) and incubated for approximately 2 minutes. FACS analysis was then performed as previously described (44).

### Western blot

Cell lysate preparation and immunoblot analysis were performed as described (57, 59). Briefly, cells or islets were lysed in cell lysis buffer containing 1mM sodium pyrophosphate, 10mM β-glycerol phosphate, 10mM sodium fluoride, 0.5mM sodium orthovanadate, 1 μM microcystin, and protease inhibitor cocktail set II (Calbiochem) (36, 37). Supernatants of the lysates were subjected to SDS-PAGE (10-12% polyacrylamide gel) and then transferred to PVDF membranes. Antibodies used: Phopho-AKT (Ser473) (CST #4060), Phospho-p70 S6 Kinase (Thr389) (CST#9205), Cleaved Caspase-3 (Asp175) (CST#9661), Cyclin D1 (Santa Cruz, sc-8396), Cyclin D2 (Santa Cruz, sc-593). ECL Secondary Mouse and Rabbit HRP antibodies used were from GE Healthcare.

### RNA sequencing and Data Analysis

Six mouse islet preparations (3 in Normal chow group and 3 in HFD group, on diet for 4 month) were sequenced and data was analyzed using Partek Flow and IPA (Ingenuity Pathway Analysis). In brief, RNeasy Mini Kit (Qiagen, Cat# 74104) was used to isolate total RNA from mice islets. RNA quality was tested using Agilent Bioanalyzer and the RNA integrity number (RIN) values for all the samples were >7.5. We utilized USC NGS core services to convert mRNA to cDNA libraries which was further sequenced using Illumina NextSeq500-25 million reads per sample.

### Statistical Analysis

The data are presented as means ± the standard error of the mean (SEM). Differences between HFD or PA treated groups vs. controls were analyzed by Student’s t test, with two-tailed p values less than 0.05 considered statistically significant. All tissue culture experiments were performed at least three times.

## Acknowledgement

This work was supported by NIH grants R01DK084241 (BLS). Dr. Stiles also acknowledges support from NCI (R01CA154986). We also acknowledge support from the USC center for Liver Disease (P30DK48522). The funders had no role in study design, data collection and interpretation, or the decision to submit the work for publication.

## Author Contribution

R.A. wrote the manuscript and conducted experiments. Z. P., N. Z, J. S., L. H., C.C., J.C., and A. D. conducted experiments. B. L. S. directed the project, wrote and edited the manuscript.

